# Backstepping Based Automatically Controlled Angiogenic Inhibition Therapy

**DOI:** 10.1101/2020.01.13.905067

**Authors:** R. Ozgur Doruk

## Abstract

In this work, we present an automatically controlled angiogenic inhibition therapy where the variation profile of the inhibitory agent is generated by a control law that is derived using a back-stepping based control methodology. The angiogenic inhibition is described by a second order model representing the dynamics of tumor and supporting vasculature volumes. Backstepping control recursively stabilizes that dynamics and generates automatic control laws that allows the tumor volume to stay at a desired value. The desired value will be kept at one thousandth of its initial value. This will lead to a very small injection at the steady state. This is important as zero injection may lead to regrow of the tumor. The results will be presented in tabular and graphical forms. Tabular results present the variation of maximum injection rate and setup time. Graphical results present the variation of tumor and supporting vasculature volumes, injection rate and the tracking error between the reference and actual tumor volumes. In addition, we will also perform a simulation to test the capability of the closed loop to accomodate the parametric uncertainties in the rate constants. The uncertainties are represented by a random deviation in the range ±10% times the nominal value of the effected parameter. The control laws will be kept the same and the simulations will be repeated by 1000 times and each result will be superimposed on the graph. The area occupied by the curves will show the relative capability of the designs.

## 1 Angiogenic Inhibition Model of Cancer Progression

In this research we will utilize the angiogenic inhibition model developed by [1] and [2]. That is a nonlinear system represented by the following differential equations:

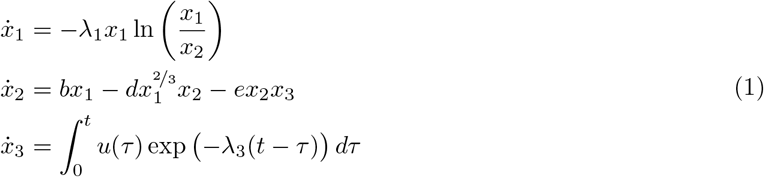

where *x*_1_ is the tumor volume in mm^3^, *x*_2_ is the supporting vasculature volume in mm^3^ (blood vessels stem from a regular vessel due to tumor angiogenesis after secretion of vascular endothelial growth factor to the site), *x*_3_ the level of inhibitory agent in the blood serum in ^mg^/kg and *u* is the inhibitor administration rate in ^mg^/kgoday. There are five parameters in (1) *θ* = [*b*, *d*, *e*, λ_1_, λ_3_] and their definitions, units and nominal values are available in **Table 1**.

**Table 1:** The numerical values of parameters in (1). These are taken from [3] which presents a mathematical model of animal Lewis lung carcinoma.

| Parameter | Unit | Value |
| --- | --- | --- |
| $b$ | $\text{day}^{-1}$ | 5.85 |
| $d$ | $\text{day}^{-1} \cdot \text{mm}^{-2}$ | 0.00873 |
| $e$ | $\text{day}^{-1} \cdot (\text{mg/kg})^{-2}$ | 0.66 |
| $\lambda_1$ | $\text{day}^{-1}$ | 0.192 |
| $\lambda_3$ | $\text{day}^{-1}$ | 1.3 |

The integral equation found in the *ẋ*_3_ equation is actually not very convenient for a control application. That is a convolution operation and defines the response of a linear system expressed by the following transfer function:

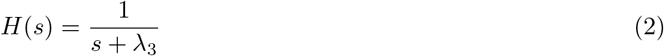

to the input *u*(*t*). The value of the parameter λ_3_ relatively closer to unity. Because of that the steady state gain of (2) is about 0.77. This corresponds to a rise time of about 1.7 days. This seems quite fast concerning a realistic problem. So one can directly replace the plasma inhibitory agent variable *x*_3_ by the inhibitory agent rate represented by *u*. This is also the case in the relevant examples presented by [4, 2]. So one can reduce the third order model in (1) to a second order one by just replacing *x*_3_ by *u* in the *ẋ*_2_ equation as:

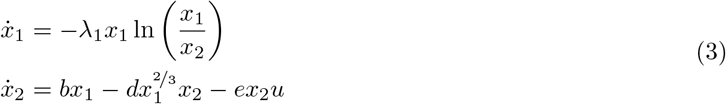

Uncontrolled evolution of the tumor volume and supporting vasculature volumes can be seen in **Figure 1**. The parameters are taken from **Table 1**.

**Figure 1:**
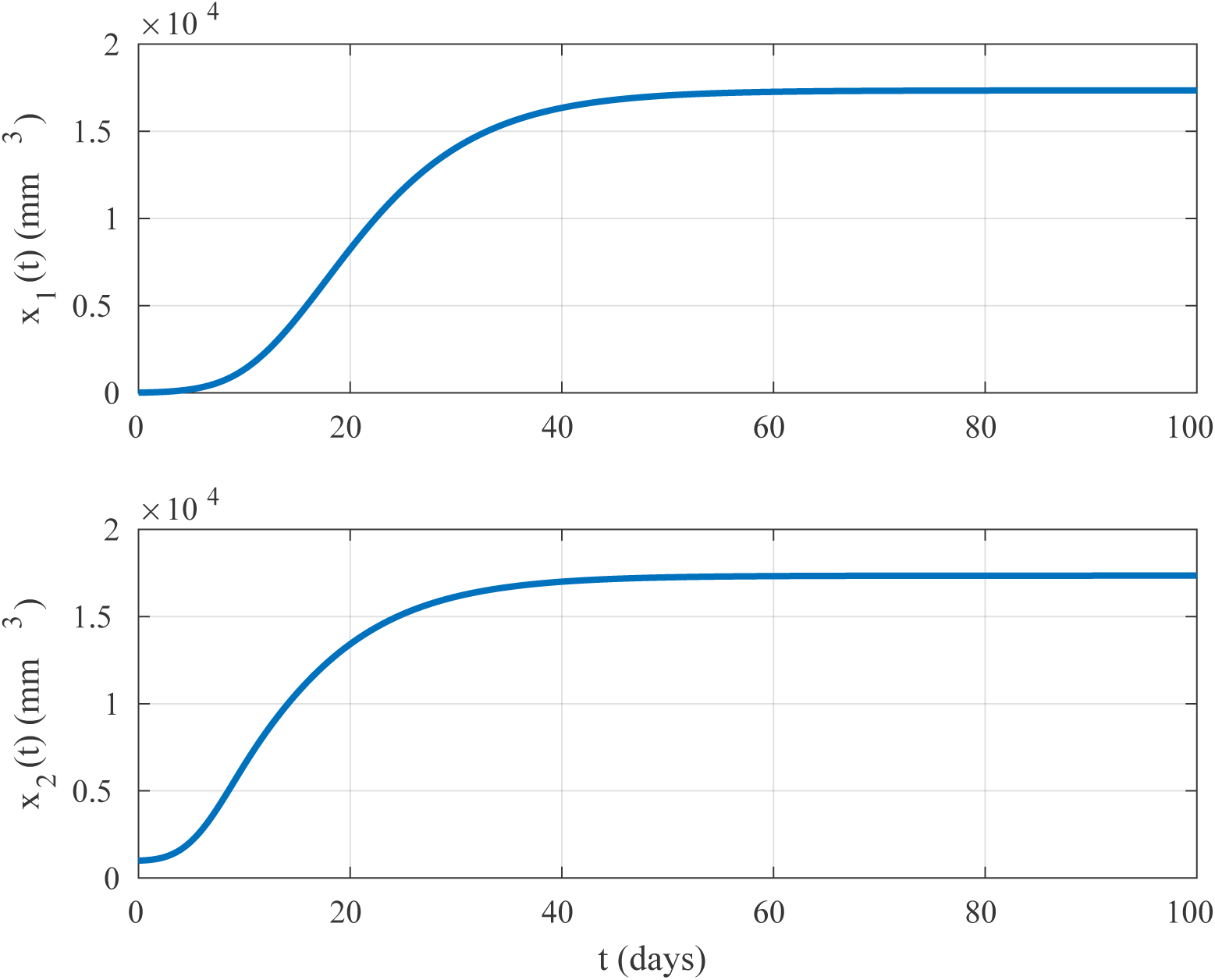
The evolution of tumor volume under no treatment conditions. The responses are obtained from (3) with the parameters in **Table 1**.

## 2 Application of Backstepping Control to Angiogenic Inhibition Problem

In this section, we will apply the famous backstepping technique [5] to derive automatically controlled inhibitory agent injection laws. One should note here that, the angiogenic inhibition model presented in (3) is a full relative degree model. This means that one can allow the backstepping based approach without the risk of an uncontrollable remaining dynamics.

### 2.1 Step 1: Stabilization of the Tumor Volume

Before proceeding one needs to define a tracking error *ϵ*_1_ between actual and reference tumor volumes (*x*_1_ and *r*_1_) as shown below:

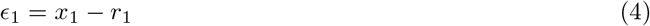

Finding its dynamics by differentiating yields:

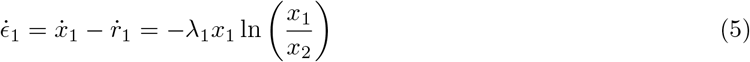

In the above we assume that the reference tumor level *r*_1_ is constant. Although we are in Step 1, one needs to define the tracking error between the supporting vasculature volume and a virtual input *r*_2_ as follows:

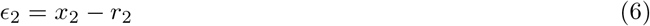

If one draws *x*_2_ as *x*_2_ = *ϵ*_2_ + *r*_2_ and substitutes into (4), we can write the following:

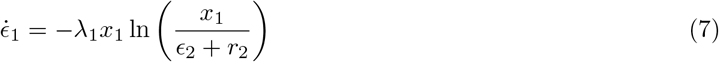

To ensure the stability, one needs to define a Control Lyapunov Function *V*_1_ as shown below:

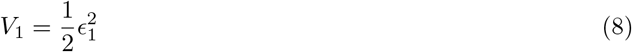

and taking its time derivative will yield:

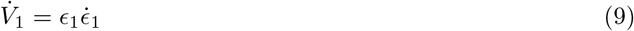

After substituting (7) to above, we will obtain:

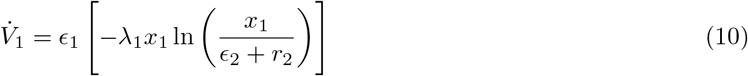

Concerning a standard backstepping application one needs to draw the virtual control variable *r*_2_ from the above equation which will yield a negative definite 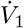. This will yield an asymptotically stable dynamics for *ϵ*_1_. One thing to note here is that backstepping techniques develop the control laws recursively. This recursiveness is provided by the Control Lyapunov Functions. In this step, this is obtained by equating (10) to a form shown by:

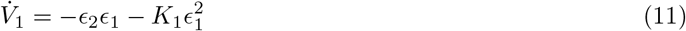

with *K*_1_ being a positive real number. In order to obtain this rate function, *r*_2_ should be derived as:

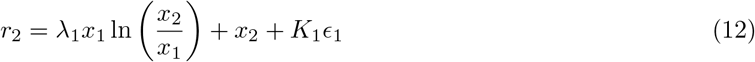

### 2.2 Step 2: Stabilization of the Supporting Vasculature Volume

One should start by taking the time derivative of the tracking error between *x*_2_ and *r*_2_ as shown in the following:

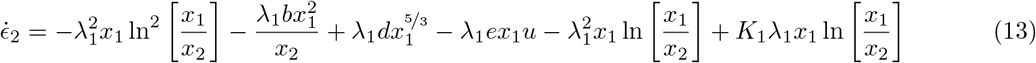

Secondly, we will need to define a Control Lyapunov Function for this step. This should be defined as an additive to that of the first one.

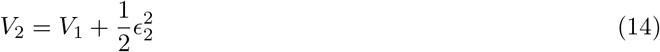

and its rate of change is:

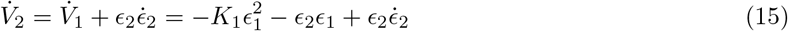

Substituting from (13) yields:

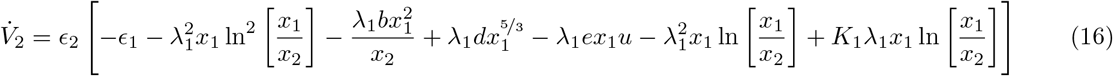

If we choose an inhibitory agent injection rate as:

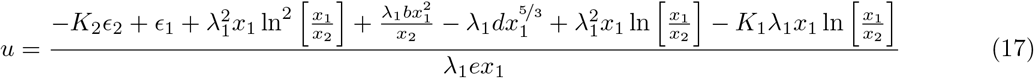

(16) becomes:

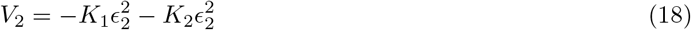

with *K*_2_ being a positive real number. (18) ensures the asymptotic stability of the closed loop.

## 3 Results

In this section we will present an example application together with its simulation results. First of all, we will present a scenario for the simulation environment and secondly we will present the results for different controller configurations.

### 3.1 Information on Simulations

One can see the information related to the initial tumor and supporting vasculature volumes and targeted treatment duration in days in **Table 2**.

**Table 2:** Problem scenario and simulation conditions

| Table 2: Problem scenario and simulation conditions |  |  |  |
| --- | --- | --- | --- |
| Scenario Parameter | Symbol | Value | Unit |
| Treatment Duration | $T_f$ | 50 | Days |
| Initial Tumor Volume | $x_1(0)$ | 2000 | mm<sup>3</sup> |
| Initial Vasculature Volume | $x_2(0)$ | 250 | mm<sup>3</sup> |
| Reference (steady state) tumor volume | $r_1$ | $\frac{1}{1000}x_1(0)$ | mm<sup>3</sup> |

The purpose of the controlled simulations is to reduce the tumor volume *x*_1_ to 1% of its initial volume *x*_1_(0) in **Table 2** in a finite duration. The duration required to achieve this goal is called as <u>setup time</u>. In the control application there is a constant reference tumor volume *r*_1_. This will be chosen as one thousandth of the initial volume i.e. *r*_1_ = 2.5 mm^3^.

### 3.2 Simulation Results: No Uncertainties Exist

A typical variation of the tumor volume *x*_1_(*t*), supporting vasculature volume *x*_2_(*t*) and injection rate of inhibitory agent *u*(*t*) can be seen in **Figure 2**.

**Figure 2:**
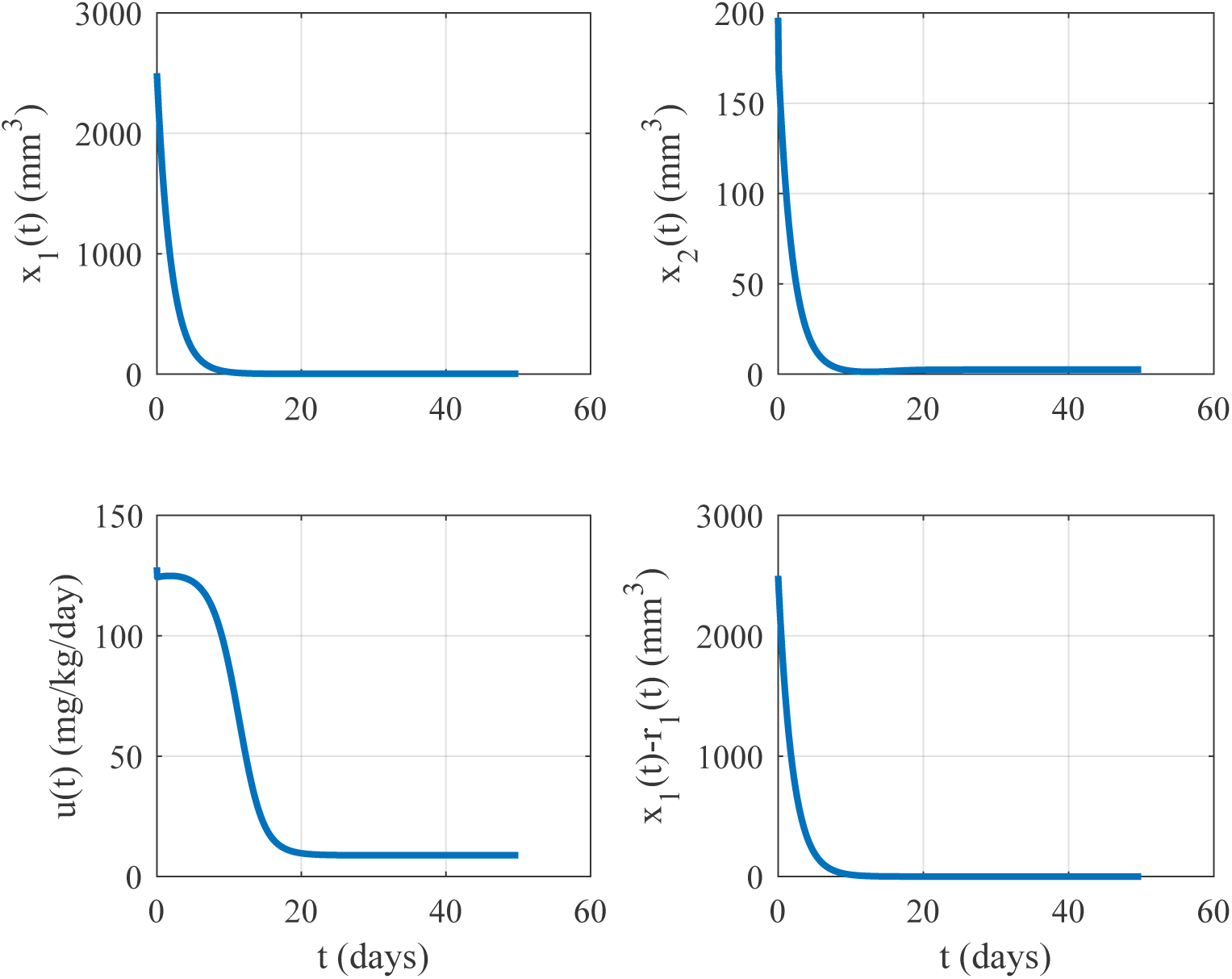
The variation of tumor volume *x*_1_(*t*), supporting vasculature volume *x*_2_(*t*) and injection rate of inhibitory agent *u*(*t*) when *K*_1_ = 100 and *K*_2_ = 0.5.

In addition to 2, one is eligible to see the results associated with maximum required injection and setup times in **Table 3**. Note that some rows in this table depicts pretty unapplicable cases (i.e. instability or very large injection rates).

**Table 3:** Maximum value of inhibitory injection rate and setup times of backstepping based treatment procedures. Here max(*u*) is the maximum inhibitory agent injection rate and *T_s_* is the setup time in days.

| $K_1$ | $K_2$ | $\max(u)$ | $T_s$ |
| --- | --- | --- | --- |
| 2.0000 | 0.0500 | 1249.3485 | 5.5320 |
| 2.0000 | 0.1000 | 2789.3545 | 5.0430 |
| 2.0000 | 0.2000 | 44372.6944 | 4.3100 |
| 2.0000 | 0.2500 | 4033991.5434 | 4.0360 |
| 5.0000 | 0.0500 | 101.4783 | 17.8450 |
| 5.0000 | 0.1000 | 103.2578 | 14.9000 |
| 5.0000 | 0.2000 | 106.8169 | 11.2200 |
| 5.0000 | 0.5000 | 400.8060 | 6.4910 |
| 5.0000 | 0.9000 | 3584.5894 | 4.2040 |
| 5.0000 | 1.5000 | 95733.1915 | 2.8020 |
| 5.0000 | 2.5000 | 19222732.9454 | 1.8790 |
| 5.0000 | 5.0000 | 137296569206.2600 | 1.2380 |
| 10.0000 | 0.0100 | 81.3145 | 42.0260 |
| 10.0000 | 0.0200 | 82.0646 | 38.5310 |
| 10.0000 | 0.0500 | 84.3149 | 30.8440 |
| 10.0000 | 0.1000 | 88.0653 | 23.1570 |
| 10.0000 | 0.2000 | 95.5661 | 15.4700 |
| 10.0000 | 0.5000 | 206.7063 | 7.7840 |
| 10.0000 | 1.0000 | 2866.5315 | 4.2920 |
| 10.0000 | 2.0000 | 538543.5230 | 2.3000 |
| 20.0000 | 0.0002 | 42.3266 | 93.1330 |
| 20.0000 | 0.0020 | 42.6035 | 89.9110 |
| 20.0000 | 0.0100 | 43.8343 | 77.9290 |
| 20.0000 | 0.0200 | 45.3727 | 66.8030 |
| 20.0000 | 0.0500 | 49.9880 | 46.7770 |
| 20.0000 | 0.1000 | 57.6802 | 31.2000 |
| 20.0000 | 0.2000 | 73.0645 | 18.7390 |
| 20.0000 | 0.5000 | 155.0376 | 8.5440 |
| 20.0000 | 1.0000 | 2113.3121 | 4.4980 |
| 20.0000 | 2.0000 | 385414.3820 | 2.3270 |
| 20.0000 | 4.0000 | 12287655310.0452 | 1.2020 |
| 50.0000 | 0.0020 | 9.7487 | 213.5450 |
| 50.0000 | 0.0100 | 10.1500 | 156.6040 |
| 50.0000 | 0.0200 | 10.6819 | 117.4580 |
| 50.0000 | 0.0500 | 12.4766 | 67.1270 |
| 50.0000 | 0.1000 | 16.2033 | 39.1650 |
| 50.0000 | 0.2000 | 27.3737 | 21.3710 |
| 50.0000 | 0.5000 | 131.6975 | 9.0520 |
| 50.0000 | 1.0000 | 1789.5742 | 4.6230 |
| 50.0000 | 2.0000 | 325013.6384 | 2.3420 |
| 50.0000 | 4.0000 | 10618811896.1328 | 1.1810 |
| 100.0000 | 0.0020 | 9.2806 | 391.9980 |
| 100.0000 | 0.0100 | 9.6517 | 235.2020 |
| 100.0000 | 0.0200 | 10.1496 | 156.8040 |
| 100.0000 | 0.0500 | 11.8434 | 78.4060 |
| 100.0000 | 0.1000 | 15.3737 | 42.7710 |
| 100.0000 | 0.5000 | 128.4096 | 9.2310 |
| 100.0000 | 1.0000 | 1696.9747 | 4.6640 |
| 100.0000 | 2.0000 | 308350.3896 | 2.3460 |
| 100.0000 | 2.2500 | 1131789.4217 | 2.0860 |

### 3.3 Simulation Results: Simulations Under Existence of Uncertainties

In this section, we will present simulations under the existence of 10% uncertainty in the rate constants (i.e. [*b*, *d*, *e*, λ_1_]). The uncertainties are incormporated into the parameters by introducing random deviations to those parameters. This deviation is obtained by adding a uniformly distributed random number in the following fashion:

For example for parameter *b*, the uncertainty is incorporated as 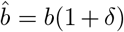 where *δ* is a random number uniformly distributed in the interval [—0,1,0,1]. This means that there is a 10% uncertainty assumed in the parameters. The deviated parameters are only used in the simulation of the angiogenic inhibition model itself. The controller stays same. In other words, controller’s design involves the nominal values of the parameters *θ* = [*b*, *d*, *e*, λ_1_].

Since we have randomly varying parameters, the simulation needs to be repeated. We repeat the simulations 1000 times and superimpose the results on the same figure. The `hold on/off` feature of the MATLAB environment draws each curve with a different color. As the number of runs advances, the curves may occupy a larger space. The larger the occupied area the larger the deviations.

The simulations are performed with the configurations presented in **Table 4**. If the results are not therapeutically feasible under 10% uncertainty, the situation is also indicated. The results associated with the therapeutically feasible cases are presented in **Figures 3–16** referenced in the table. It should be noted that, unstable and cases resulting negative inhibitory agent injections are not reported graphically.

**Figure 3:**
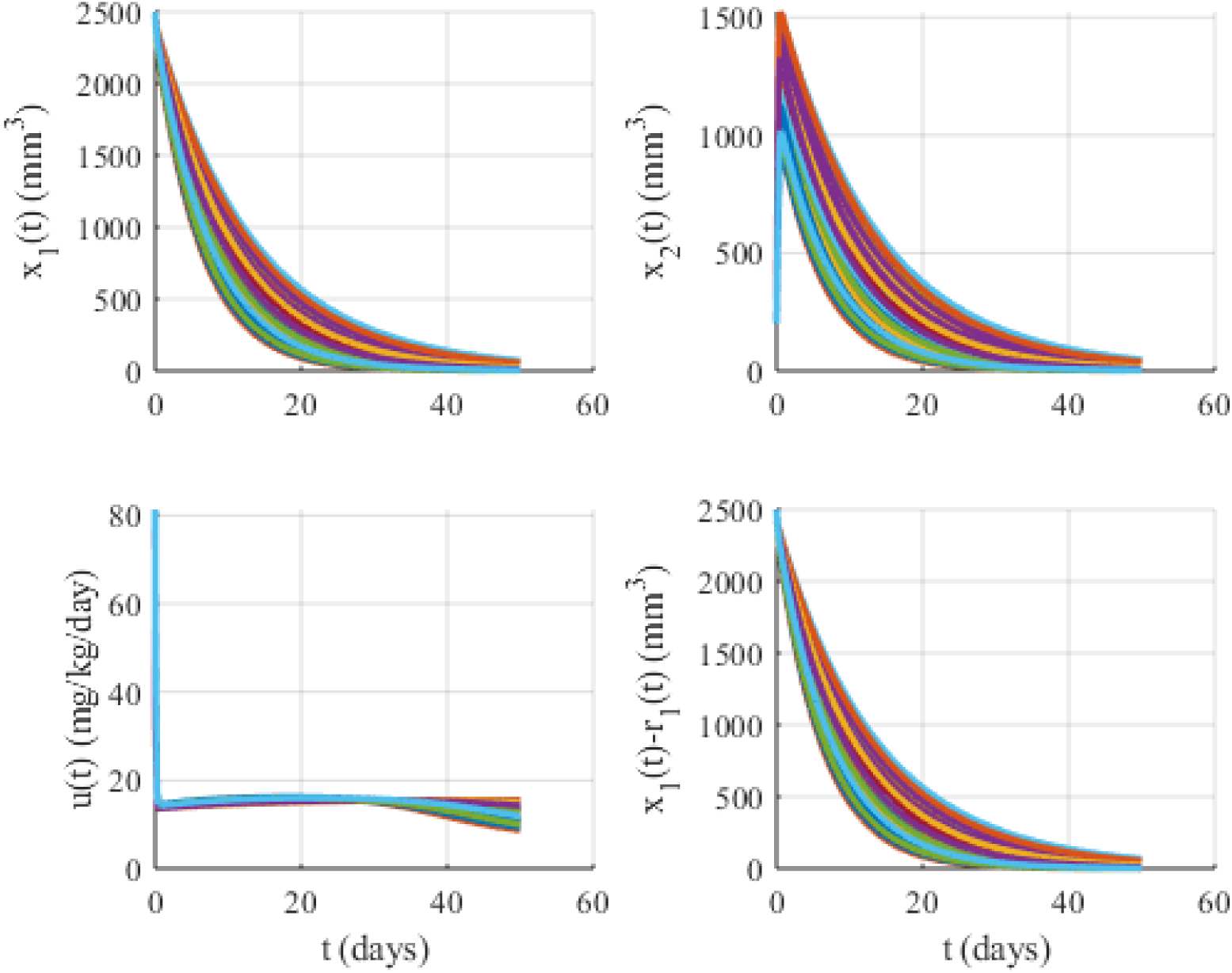
Simulations under the existence of 10% uncertainty when *K*_1_ = 10 and *K*_2_ = 0.01. *x*_1_(*t*): Tumor volume, *x*_2_(*t*): Supporting vasculature volume, *u*(*t*): Inhibitory agent rate.

**Figure 4:**
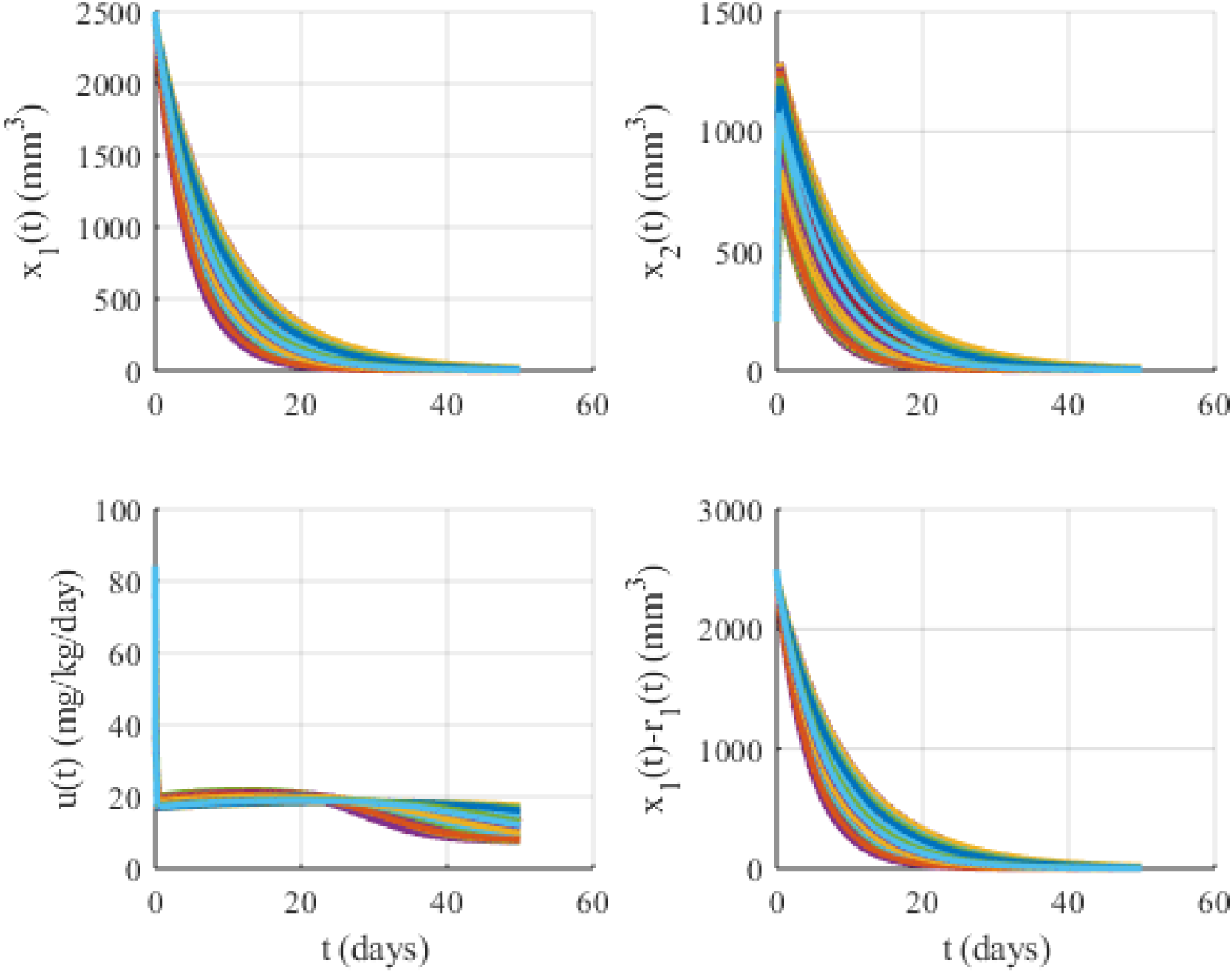
Simulations under the existence of 10% uncertainty when *K*_1_ = 10 and *K*_2_ = 0.05. *x*_1_(*t*): Tumor volume, *x*_2_(*t*): Supporting vasculature volume, *u*(*t*): Inhibitory agent rate.

**Figure 5:**
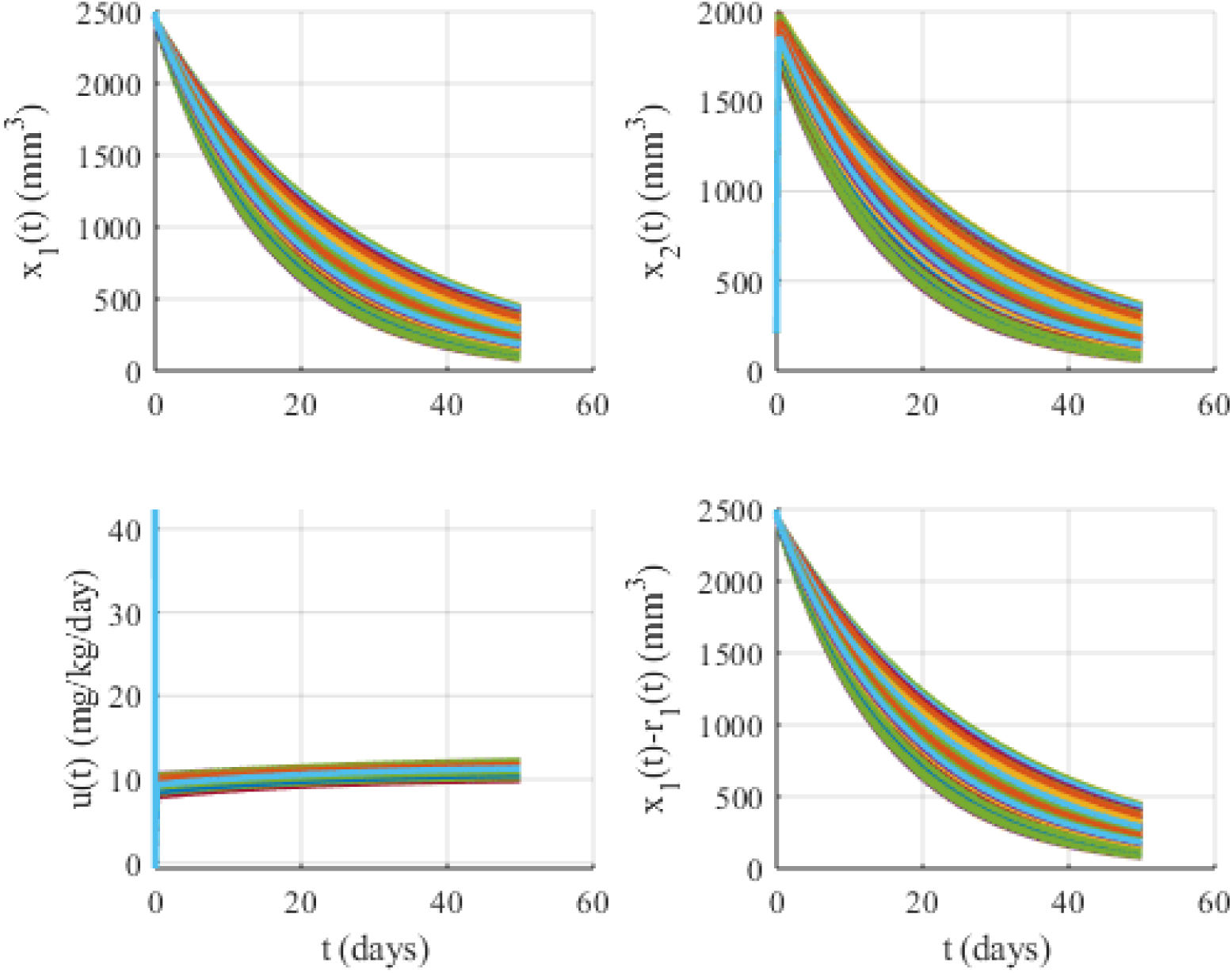
Simulations under the existence of 10% uncertainty when *K*_1_ = 20 and *K*_2_ = 0.0002. *x*_1_(*t*): Tumor volume, *x*_2_(*t*): Supporting vasculature volume, *u*(*t*): Inhibitory agent rate.

**Figure 6:**
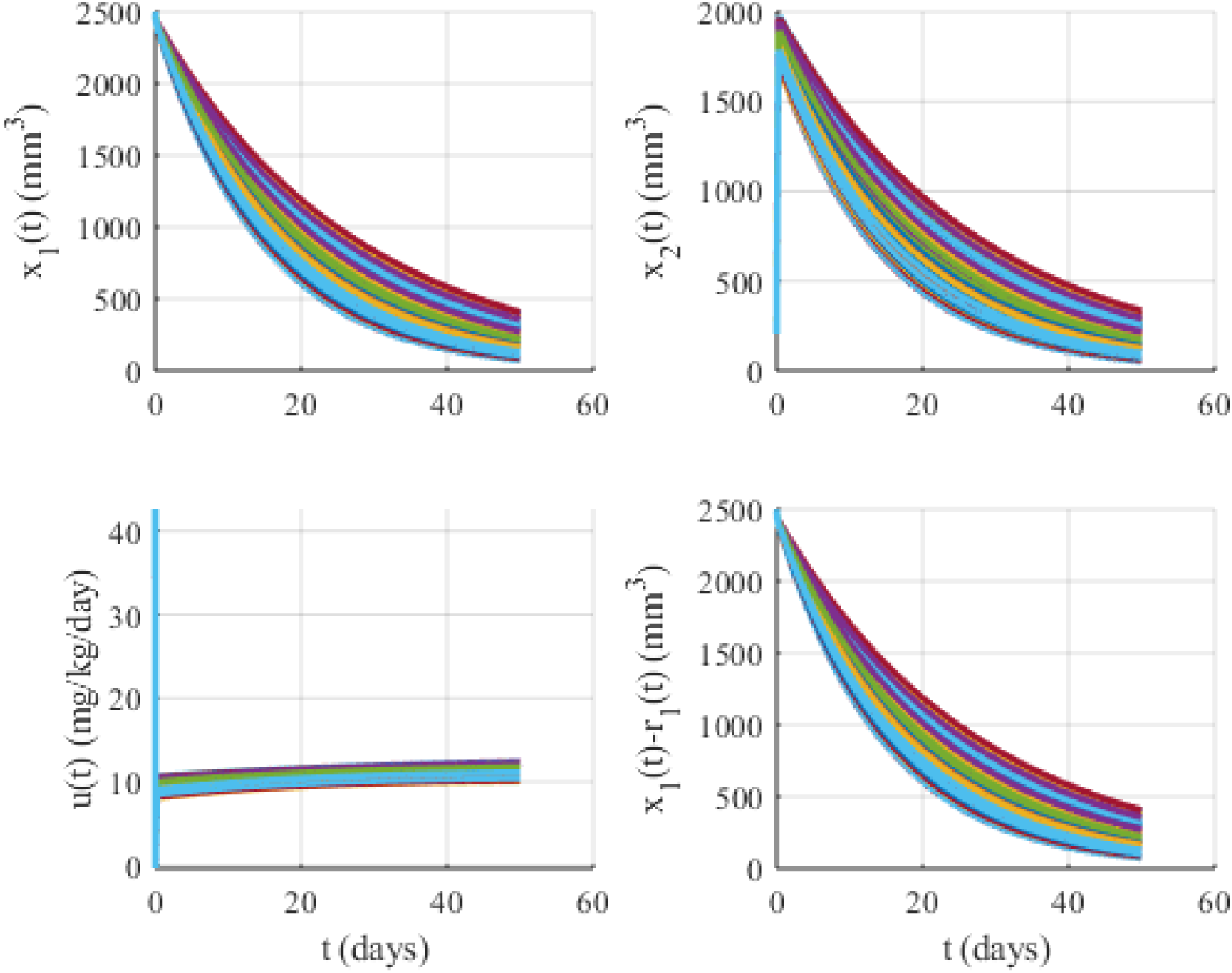
Simulations under the existence of 10% uncertainty when *K*_1_ = 20 and *K*_2_ = 0.002. *x*_1_(*t*): Tumor volume, *x*_2_(*t*): Supporting vasculature volume, *u*(*t*): Inhibitory agent rate.

**Figure 7:**
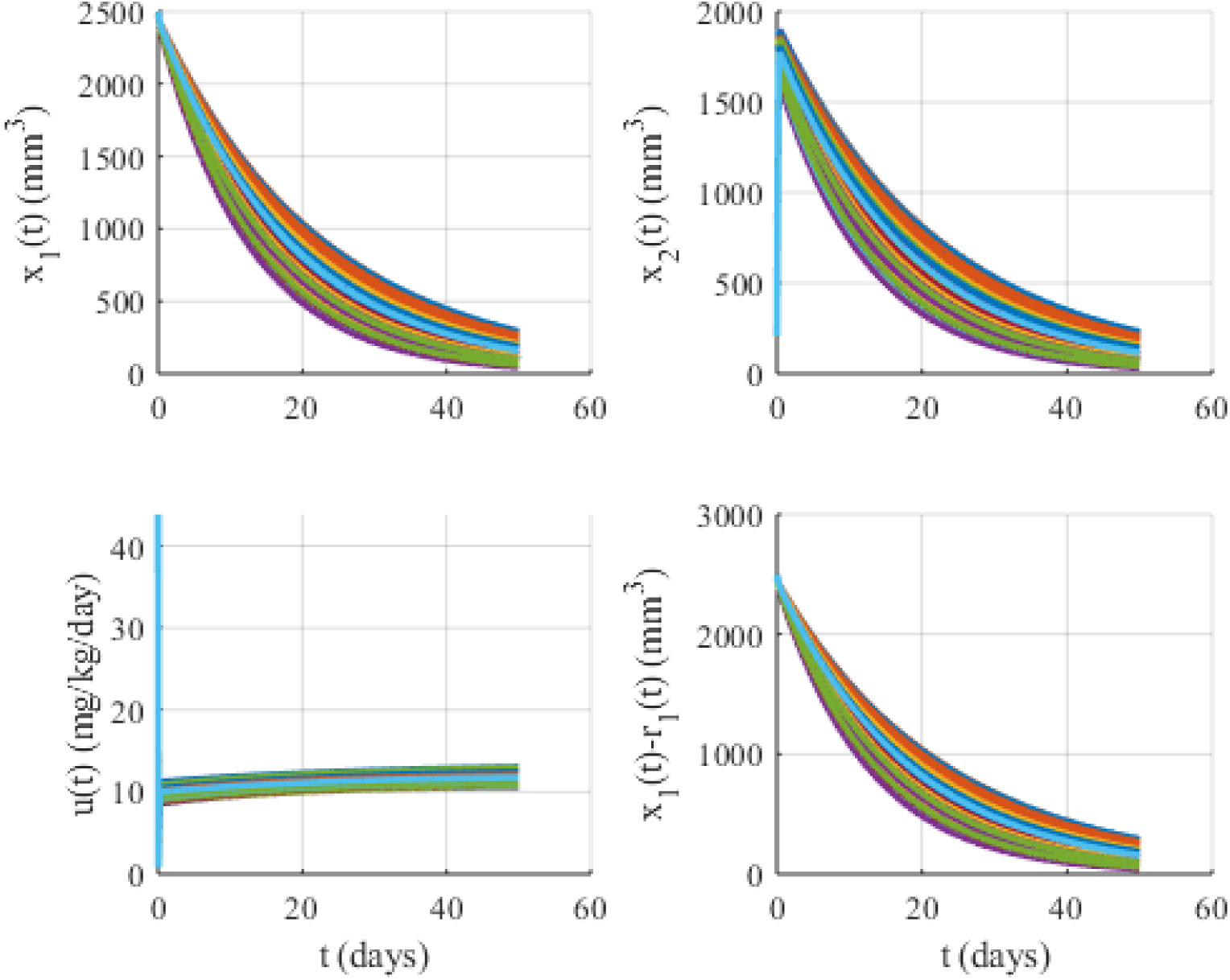
Simulations under the existence of 10% uncertainty when *K*_1_ = 20 and *K*_2_ = 0.01. *x*_1_(*t*): Tumor volume, *x*_2_(*t*): Supporting vasculature volume, *u*(*t*): Inhibitory agent rate.

**Figure 8:**
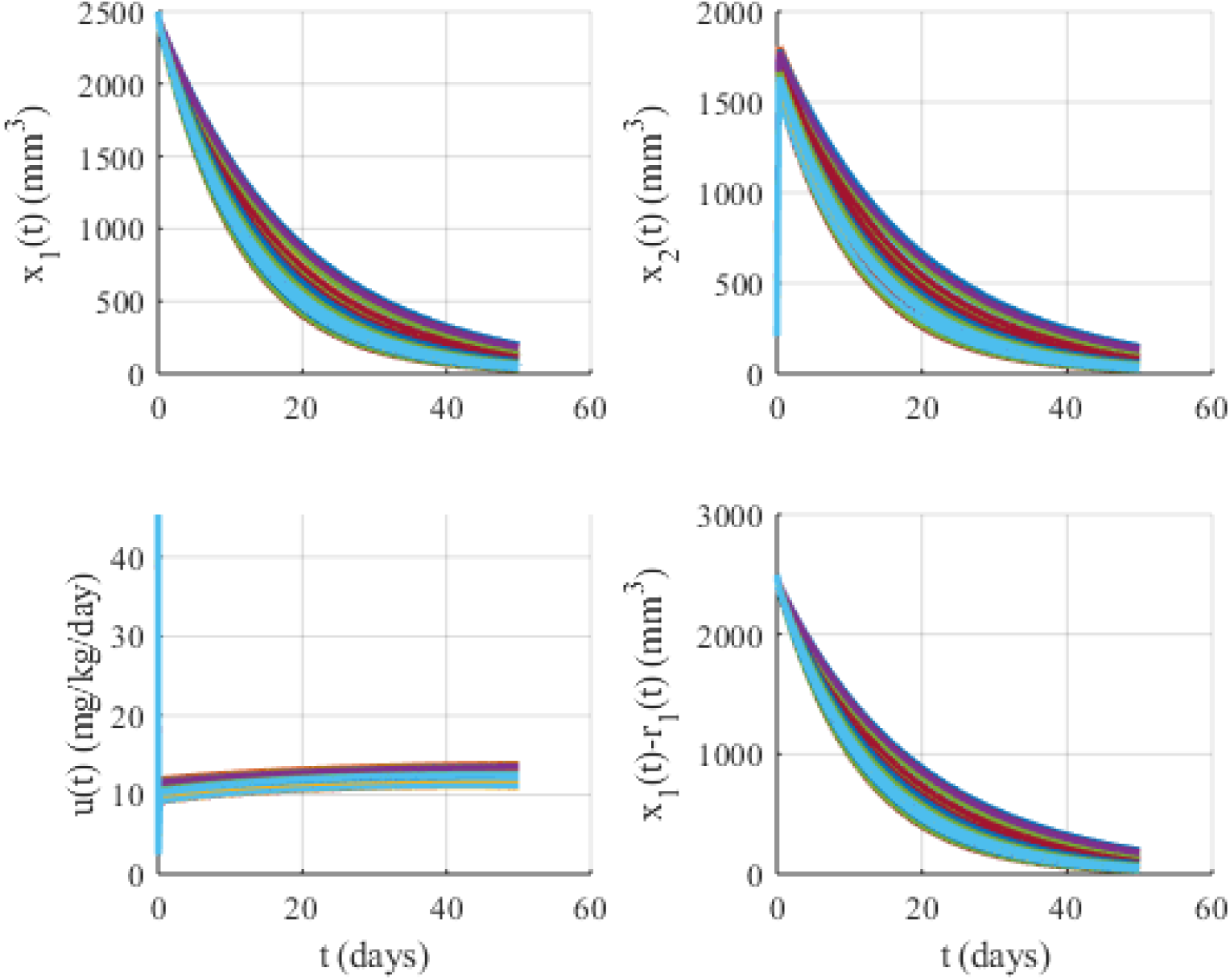
Simulations under the existence of 10% uncertainty when *K*_1_ = 20 and *K*_2_ = 0.02. *x*_1_(*t*): Tumor volume, *x*_2_(*t*): Supporting vasculature volume, *u*(*t*): Inhibitory agent rate.

**Figure 9:**
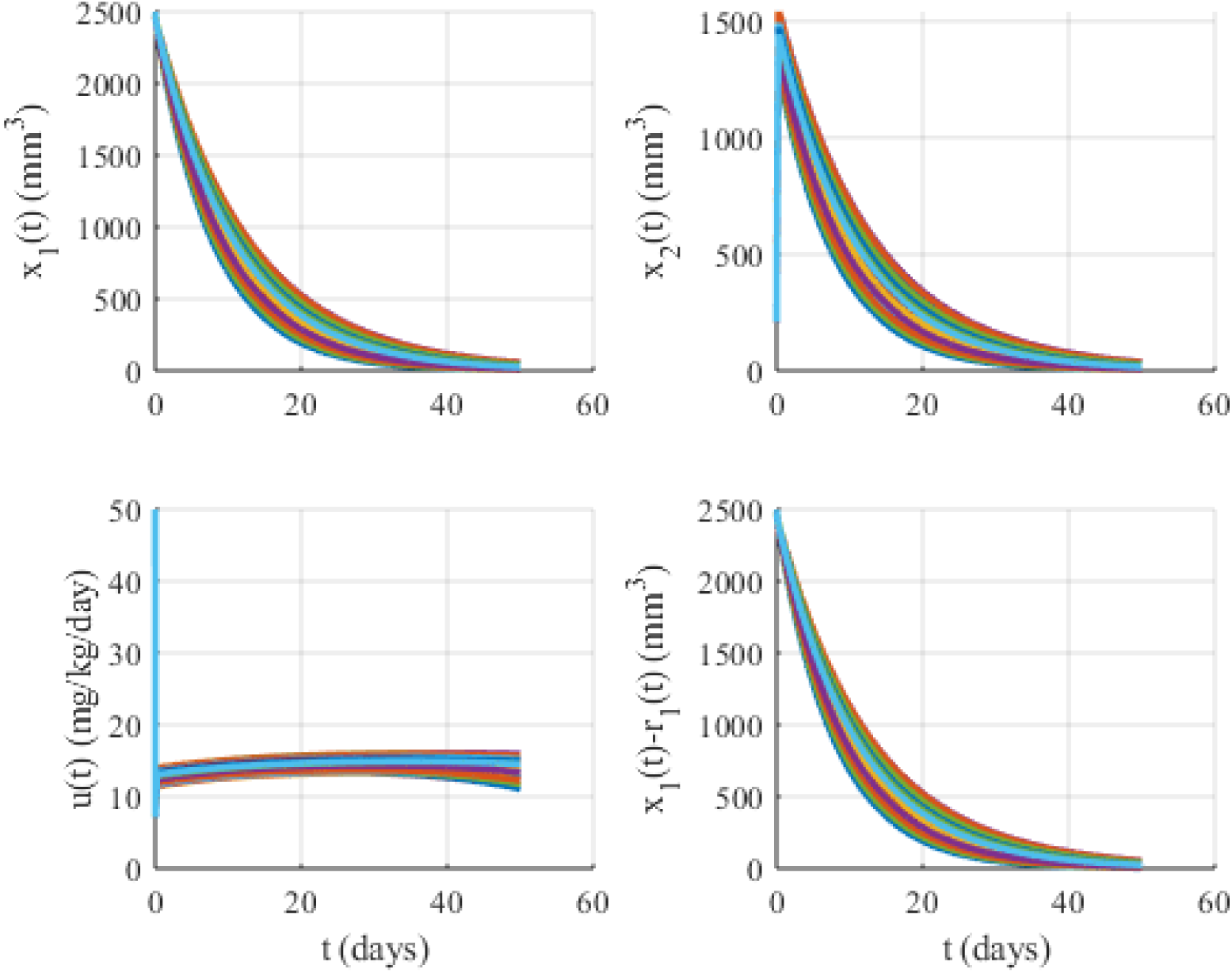
Simulations under the existence of 10% uncertainty when *K*_1_ = 20 and *K*_2_ = 0.05. *x*_1_(*t*): Tumor volume, *x*_2_(*t*): Supporting vasculature volume, *u*(*t*): Inhibitory agent rate.

**Figure 10:**
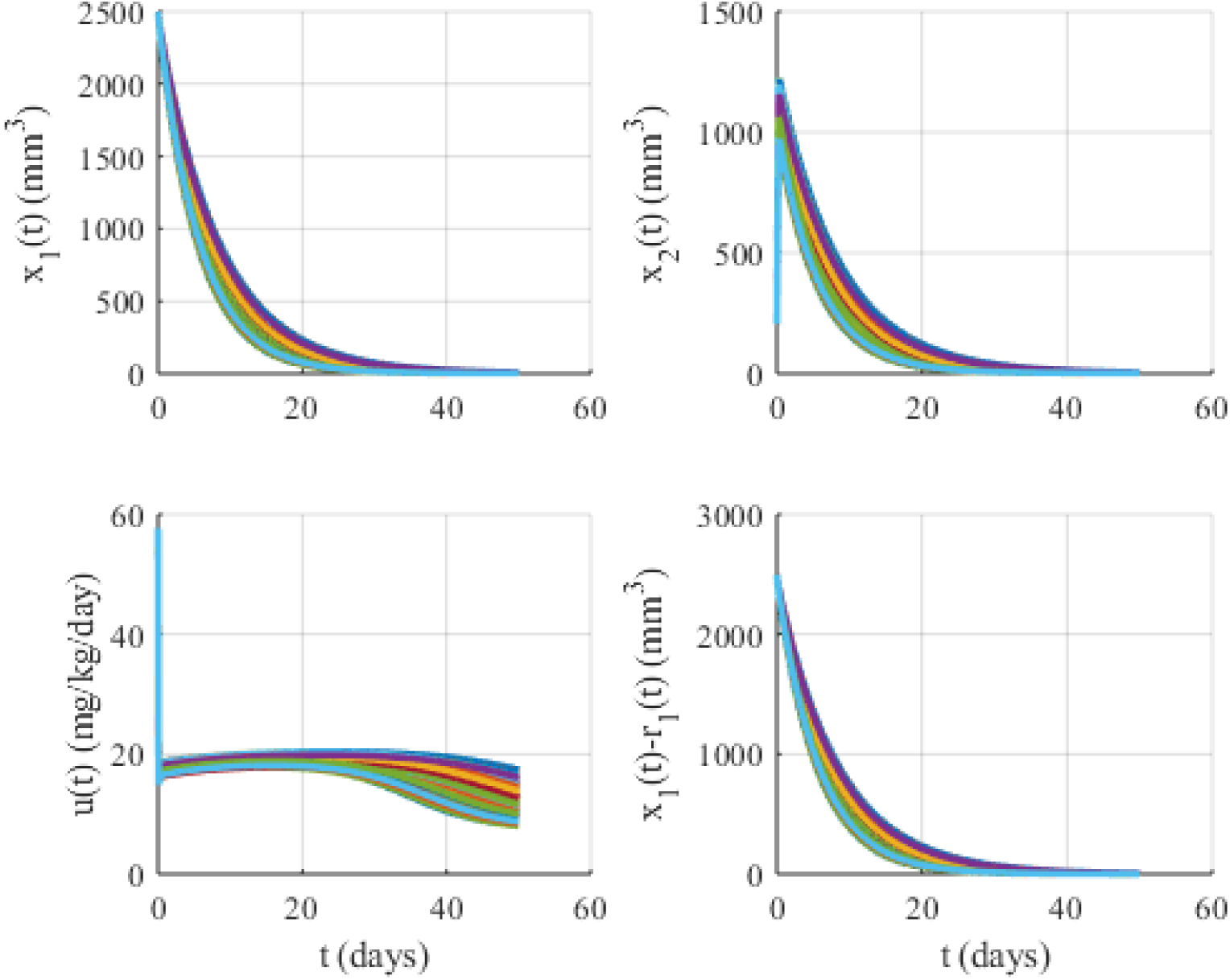
Simulations under the existence of 10% uncertainty when *K*_1_ = 20 and *K*_2_ =0.1. *x*_1_(*t*): Tumor volume, *x*_2_(*t*): Supporting vasculature volume, *u*(*t*): Inhibitory agent rate.

**Figure 11:**
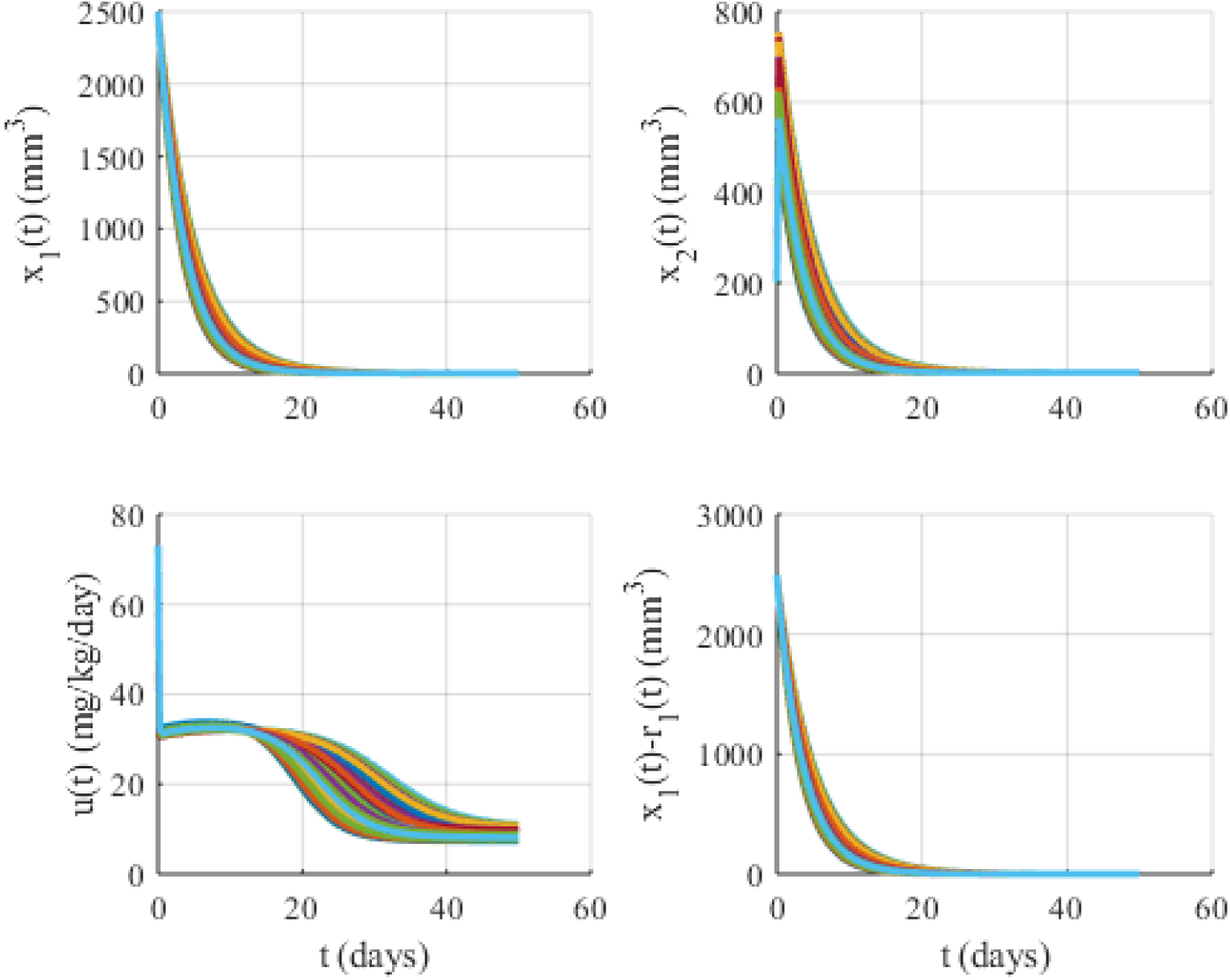
Simulations under the existence of 10% uncertainty when *K*_1_ = 20 and *K*_2_ = 0.2. *x*_1_(*t*): Tumor volume, *x*_2_(*t*): Supporting vasculature volume, *u*(*t*): Inhibitory agent rate.

**Figure 12:**
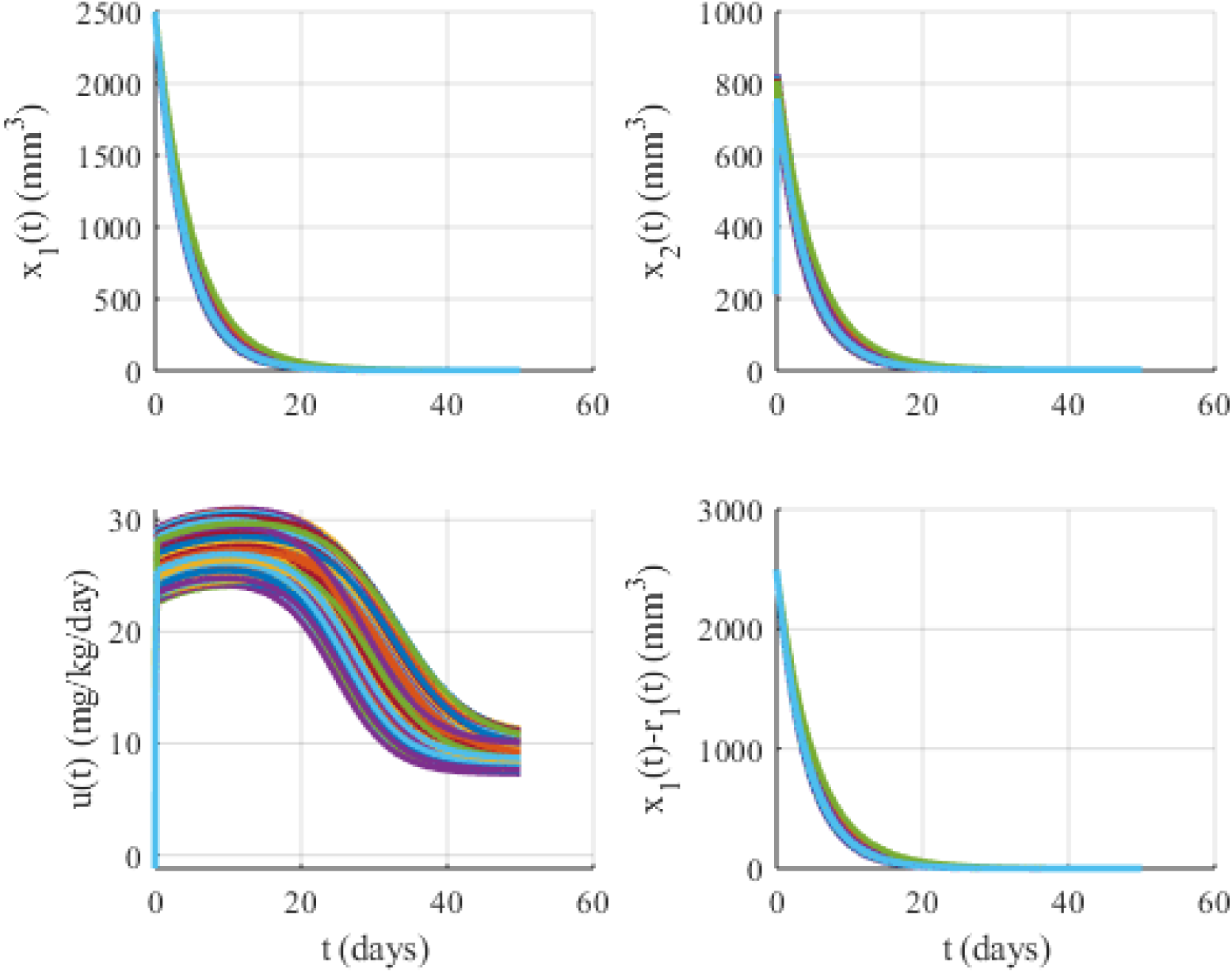
Simulations under the existence of 10% uncertainty when *K*_1_ = 50 and *K*_2_ = 0.2. *x*_1_(*t*): Tumor volume, *x*_2_(*t*): Supporting vasculature volume, *u*(*t*): Inhibitory agent rate.

**Figure 13:**
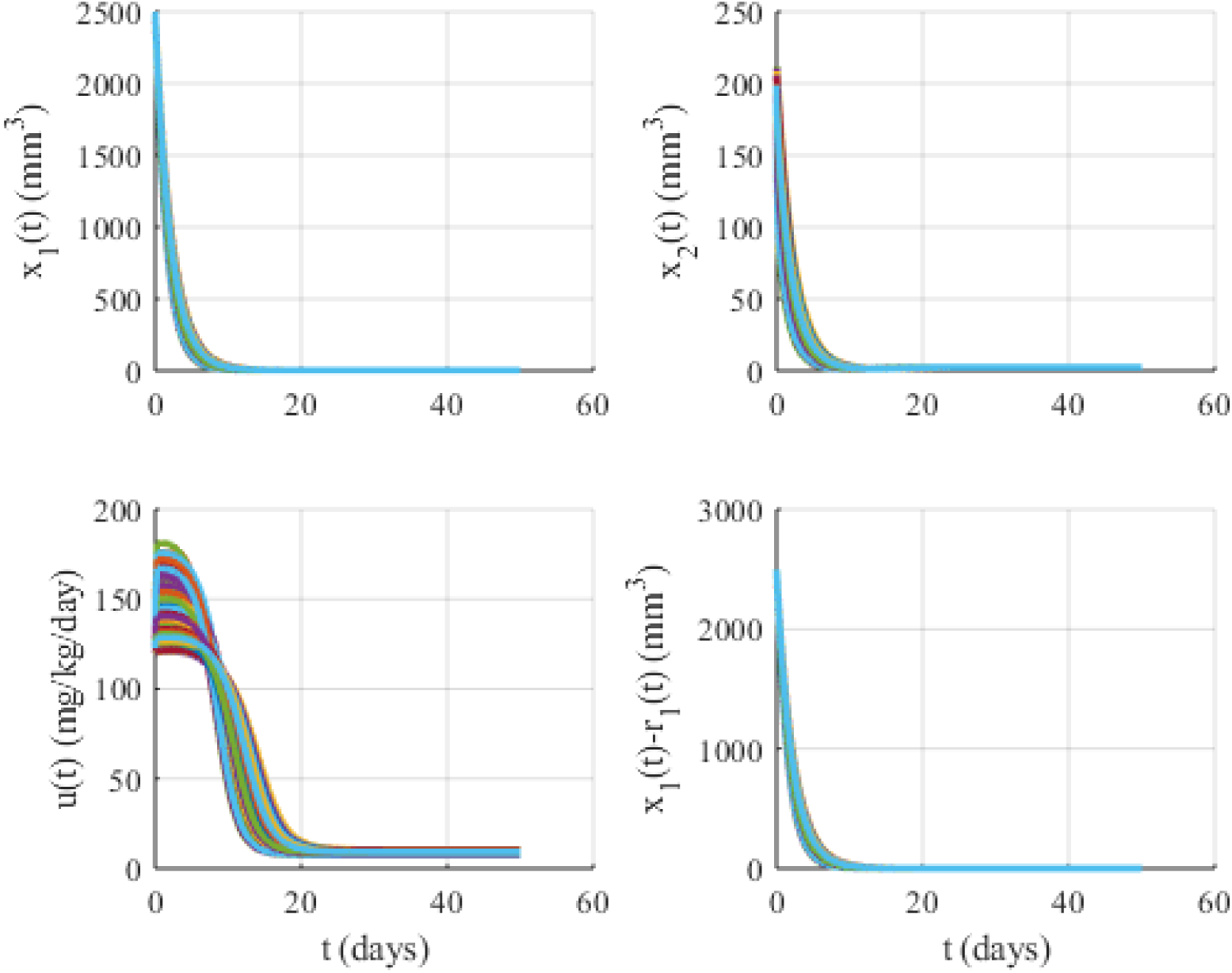
Simulations under the existence of 10% uncertainty when *K*_1_ = 50 and *K*_2_ = 0.5. *x*_1_(*t*): Tumor volume, *x*_2_(*t*): Supporting vasculature volume, *u*(*t*): Inhibitory agent rate.

**Figure 14:**
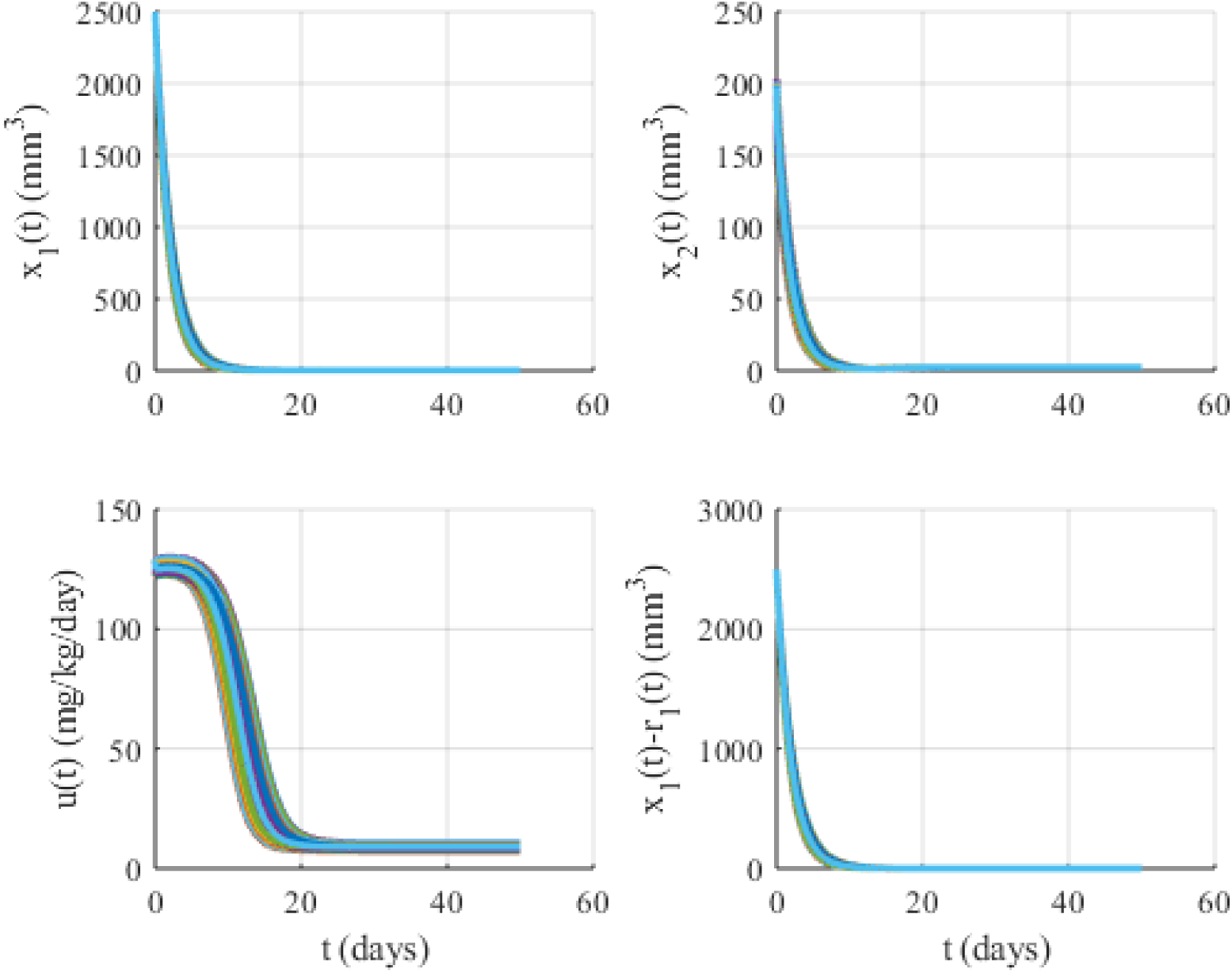
Simulations under the existence of 10% uncertainty when *K*_1_ = 100 and *K*_2_ = 0.5. *x*_1_(*t*): Tumor volume, *x*_2_(*t*): Supporting vasculature volume, *u*(*t*): Inhibitory agent rate.

**Figure 15:**
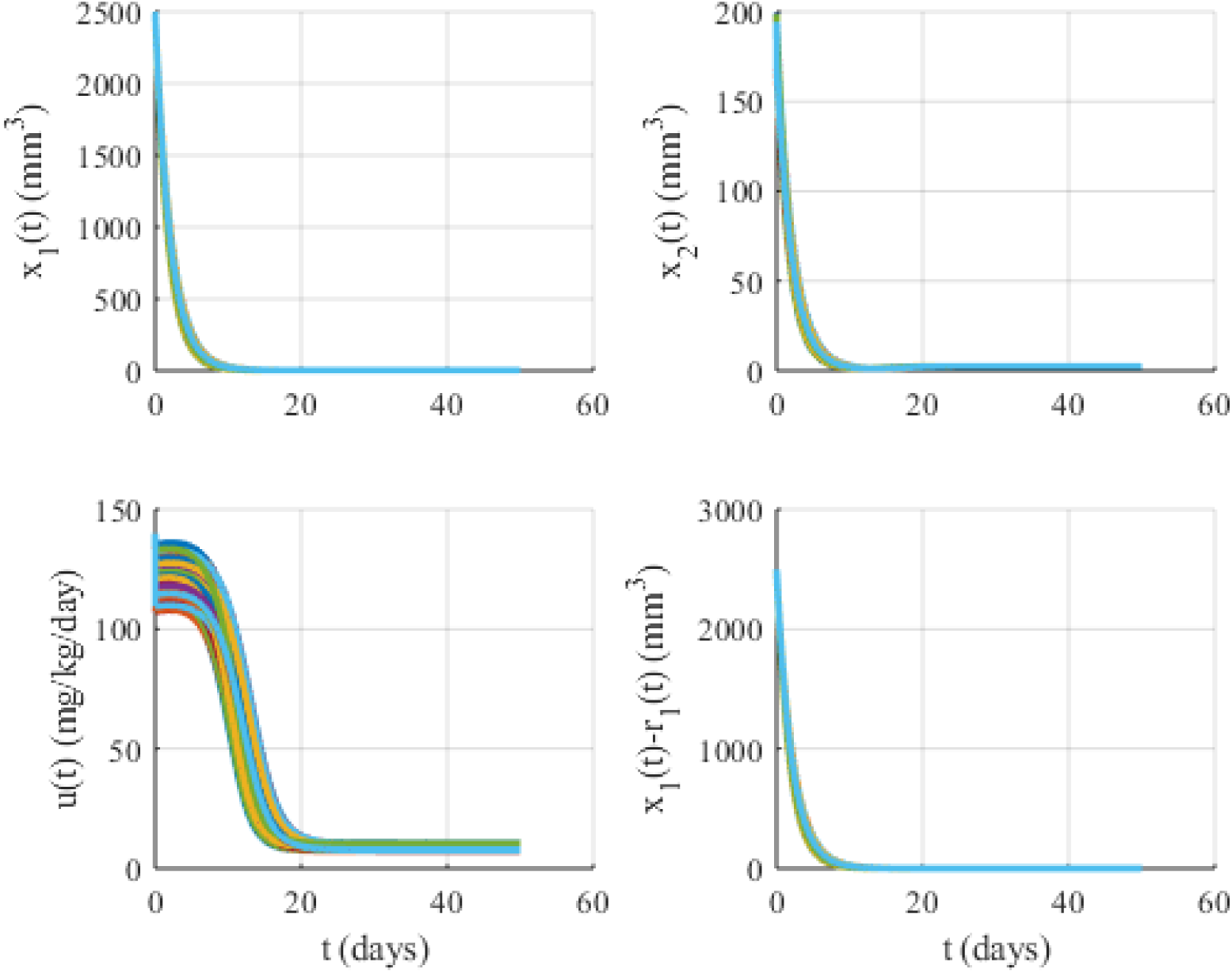
Simulations under the existence of 10% uncertainty when *K*_1_ = 200 and *K*_2_ = 0.5. *x*_1_(*t*): Tumor volume, *x*_2_(*t*): Supporting vasculature volume, *u*(*t*): Inhibitory agent rate.

**Figure 16:**
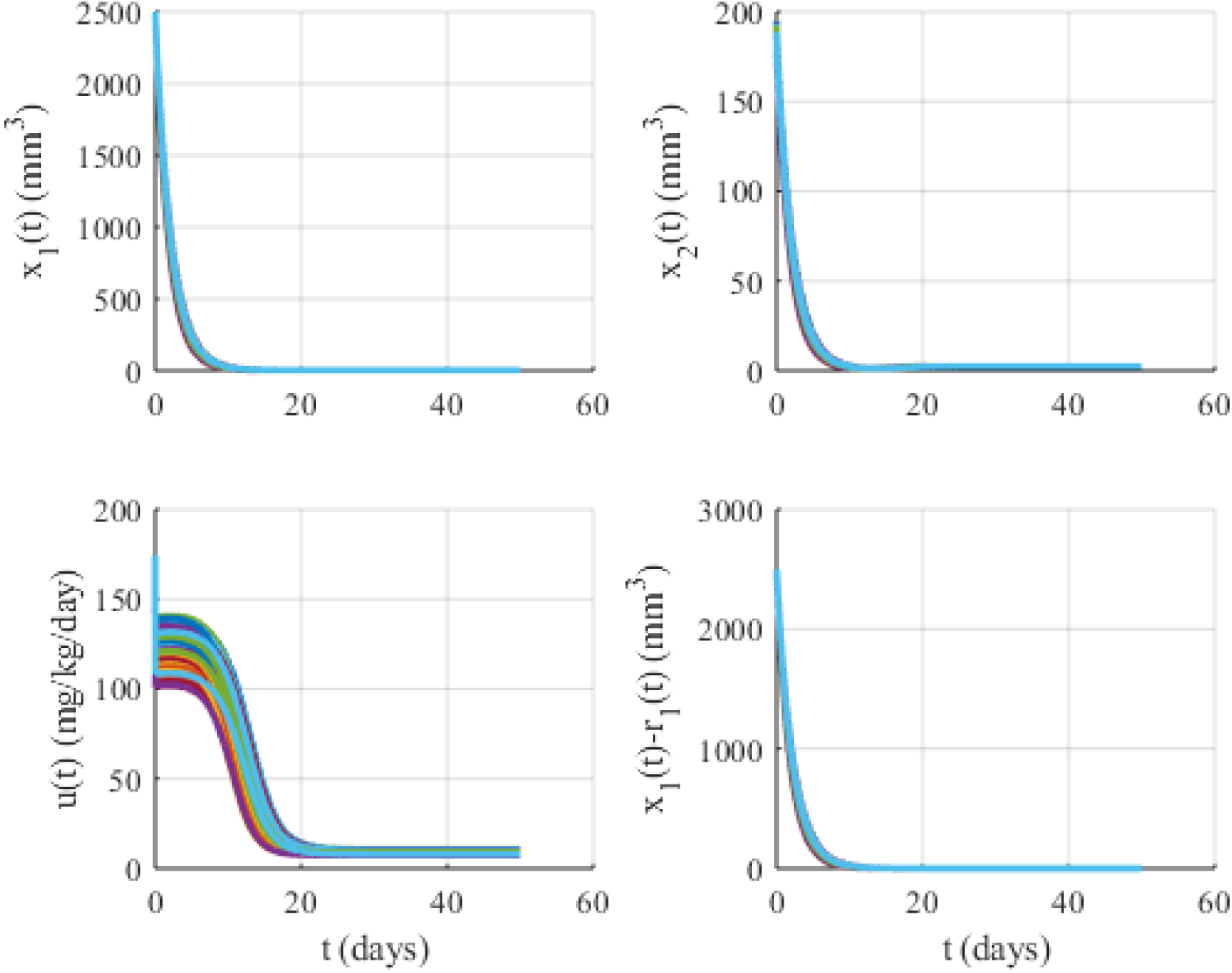
Simulations under the existence of 10% uncertainty when *K*_1_ = 500 and *K*_2_ = 0.5. *x*_1_(*t*): Tumor volume, *x*_2_(*t*): Supporting vasculature volume, *u*(*t*): Inhibitory agent rate.

**Table 4:** The configuration of the sliding surface parameters for the simulations under 10% uncertainty in the physiological rate constant parameters *θ* = [*b*, *d*, *e*, λ_1_]. If the results are not therapeutically feasible under 10% uncertainty, the situation is indicated in the observation column. Note: When the gains are swapped (i.e. *K*_1_ ⇔ *K*_2_) similar results are obtained.

| $K_1$ | $K_2$ | Observations |
| --- | --- | --- |
| 1 | 0.05 | UNSTABLE |
| 1 | 0.02 | UNSTABLE |
| 1 | 0.1 | UNSTABLE |
| 0.5 | 0.02 | UNSTABLE |
| 0.5 | 0.05 | UNSTABLE |
| 2 | 0.1 | UNSTABLE |
| 2 | 0.05 | UNSTABLE |
| 2 | 0.2 | UNSTABLE |
| 2 | 0.4 | UNSTABLE |
| 5 | 0.1 | UNSTABLE |
| 5 | 0.2 | UNSTABLE |
| 5 | 0.5 | UNSTABLE |
| 5 | 0.9 | UNSTABLE |
| 10 | 0.01 | Sudden decrease in the inhibitory agent administration rate $u(t)$.<br>Sudden jumps in the vasculature volume $x_2(t)$ |
| 10 | 0.05 | Sudden decrease in the inhibitory agent administration rate $u(t)$.<br>Sudden jumps in the vasculature volume $x_2(t)$ |
| 10 | 0.1 | Sudden decrease in the inhibitory agent administration rate $u(t)$.<br>Sudden jumps in the vasculature volume $x_2(t)$ |
| 10 | 0.2 | UNSTABLE |
| 10 | 0.5 | UNSTABLE |
| 10 | 1 | UNSTABLE |
| 10 | 2 | UNSTABLE |
| 20 | 0.0002 | Sudden decrease in the inhibitory agent administration rate $u(t)$.<br>Sudden jumps in the vasculature volume $x_2(t)$<br>Very slow response |
| 20 | 0.002 | Sudden decrease in the inhibitory agent administration rate $u(t)$.<br>Sudden jumps in the vasculature volume $x_2(t)$<br>Very slow response |
| 20 | 0.01 | Sudden decrease in the inhibitory agent administration rate $u(t)$.<br>Sudden jumps in the vasculature volume $x_2(t)$<br>Very slow response |
| 20 | 0.02 | Sudden decrease in the inhibitory agent administration rate $u(t)$.<br>Sudden jumps in the vasculature volume $x_2(t)$<br>Very slow response |
| 20 | 0.05 | Sudden decrease in the inhibitory agent administration rate $u(t)$.<br>Sudden jumps in the vasculature volume $x_2(t)$ |
| 20 | 0.1 | Sudden decrease in the inhibitory agent administration rate $u(t)$.<br>Sudden jumps in the vasculature volume $x_2(t)$ |
| 20 | 0.2 | Sudden decrease in the inhibitory agent administration rate $u(t)$.<br>Sudden jumps in the vasculature volume $x_2(t)$ |
| 20 | 0.5 | UNSTABLE |
| 20 | 1 | UNSTABLE |
| 20 | 2 | UNSTABLE |
| 20 | 4 | UNSTABLE |
| 50 | 0.002 | NEGATIVE INJECTION |
| 50 | 0.01 | NEGATIVE INJECTION |
| 50 | 0.02 | NEGATIVE INJECTION |
| 50 | 0.05 | NEGATIVE INJECTION |
| 50 | 0.1 | NEGATIVE INJECTION |
| 50 | 0.2 | Sudden decrease in the inhibitory agent administration rate $u(t)$.<br>Sudden jumps in the vasculature volume $x_2(t)$<br>Relatively faster. |
| 50 | 0.5 | Sudden decrease in the inhibitory agent administration rate $u(t)$.<br>Sudden jumps in the vasculature volume $x_2(t)$<br>Smoother responses |
| 50 | 1 | UNSTABLE |
| 50 | 2 | UNSTABLE |
| 50 | 5 | UNSTABLE |
| 100 | 0.002 | NEGATIVE INJECTION |
| 100 | 0.01 | NEGATIVE INJECTION |
| 100 | 0.02 | NEGATIVE INJECTION |
| 100 | 0.05 | NEGATIVE INJECTION |
| 100 | 0.1 | NEGATIVE INJECTION |
| 100 | 0.5 | Fast and smoother responses, no jumps. |
| 100 | 1 | UNSTABLE |
| 100 | 2 | UNSTABLE |
| 100 | 5 | UNSTABLE |
| 100 | 10 | UNSTABLE |
| 200 | 0.2 | NEGATIVE INJECTION |
| 200 | 0.5 | Fast and smoother responses, no jumps.<br>Relatively larger injection rate. |
| 200 | 1 | UNSTABLE |
| 200 | 2 | UNSTABLE |
| 200 | 5 | UNSTABLE |
| 200 | 10 | UNSTABLE |
| 500 | 0.2 | NEGATIVE INJECTION |
| 500 | 0.5 | Fast responses, jumps reappear.<br>Relatively larger injection rate. |
| 500 | 1 | NEGATIVE INJECTION |
| 500 | 2 | UNSTABLE |
| 500 | 5 | UNSTABLE |

## 4 Results, Discussion and Observation

In this text, we presented an automatically controlled angiogenic inhibition problem where the control laws are derived recursively using a back-stepping based approach. The mechanism samples the tumor and supporting vasculature volumes and generates an inhibitory agent variation profile. One can have the following observations:

1. Among the cases in **Table 4**, the most feasible configuration seems the one with *K*_1_ = 100 and *K*_2_ = 0.5. The variations of the variables are relatively faster, smooth and lowest dispersion seen under various parametric deviation. The setup time seems 9.23 days and highest level of inhibitory agent injection is 128.4 ^mg^/kg·day. See **Figure 14**.
2. Most of the cases in **Table 4** yields sudden decreases in the inhibitory agent administration rate *u*(*t*) and jumps in the supporting vasculature volume *x*_2_(*t*).
3. A robust control approach with better uncertainty accommodation such as sliding mode may be preferred.

## References

[1] P. Hahnfeldt, D. Panigrahy, J. Folkman, L. Hlatky, Tumor development under angiogenic signaling: a dynamical theory of tumor growth, treatment response, and postvascular dormancy, Cancer research 59 (19) (1999) 4770–4775.

[2] A. Szeles, D. A. Drexler, J. Sapi, I. Harmati, L. Kovacs, Model-based angiogenic inhibition of tumor growth using adaptive fuzzy techniques, Periodica Polytechnica Electrical Engineering and Computer Science 58 (1) (2014) 29–36.

[3] B. Rashidi, M. Yang, P. Jiang, E. Baranov, Z. An, X. Wang, A. Moossa, R. Hoffman, A highly metastatic lewis lung carcinoma orthotopic green fluorescent protein model, Clinical & experimental metastasis 18 (1) (2000) 57–60.

[4] S. Herdjunanto, Simple control design using double feedback structure for reducing tumor size, in: Proceedings of 2016 1st International Conference on Biomedical Engineering: Empowering Biomedical Technology for Better Future, IBIOMED 2016, 2017. URL http://www.scopus.com

[5] P. Kokotovic, M. Arcak, Constructive nonlinear control: a historical perspective, Automatica 37 (5) (2001) 637–662.

